# Gene Expression Variation in Arabidopsis Embryos at Single-Nucleus Resolution

**DOI:** 10.1101/2021.03.26.437151

**Authors:** Ping Kao, Michael A. Schon, Magdalena Mosiolek, Michael D. Nodine

**Affiliations:** Gregor Mendel Institute (GMI), Austrian Academy of Sciences, Vienna Biocenter (VBC), Dr. Bohr-Gasse 3, 1030 Vienna, Austria; Laboratory of Molecular Biology, Wageningen University, Wageningen, 6708 PB, the Netherlands

**Keywords:** Arabidopsis, embryo, RNA-seq, gene expression, transcription factor, epigenetic

## Abstract

Soon after fertilization of egg and sperm, plant genomes become transcriptionally activated and drive a series of coordinated cell divisions to form the basic body plan during embryogenesis. Early embryonic cells rapidly diversify from each other, and investigation of the corresponding gene expression dynamics can help elucidate underlying cellular differentiation programs. However, current plant embryonic transcriptome datasets either lack cell-specific information or have RNA contamination from surrounding non-embryonic tissues. We have coupled fluorescence-activated nuclei sorting together with single-nucleus mRNA sequencing to construct a gene expression atlas of *Arabidopsis thaliana* early embryos at single-cell resolution. In addition to characterizing cell-specific transcriptomes, we found evidence that distinct epigenetic and transcriptional regulatory mechanisms operate across emerging embryonic cell types. These datasets and analyses, as well as the approach we devised, are expected to facilitate the discovery of molecular mechanisms underlying pattern formation in plant embryos.

**Summary statement:** A transcriptome atlas of Arabidopsis embryos constructed from single nuclei reveals cell-specific epigenetic and transcriptional regulatory features.

## INTRODUCTION

Metazoans and land plants establish their body plans during embryogenesis (Dresselhaus and Jürgens, 2021; Gerri et al., 2020), and corresponding gene regulatory mechanisms have evolved independently in these two major eukaryotic lineages to help generate the immense morphological diversity observed in nature (Bai, 2015; Clark et al., 2006; Meyerowitz, 2002). For example, in animals it has been long recognized that maternal gene products control initial pattern formation before the transition of control from the maternal to the zygotic genome (Lee et al., 2014; Tadros and Lipshitz, 2009). By contrast, transcriptional activation of the zygotic genome soon after fertilization is necessary for zygote elongation and initial divisions in *Nicotiana tabacum* (tobacco) (Zhao et al., 2011) and the model flowering plant *Arabidopsis thaliana* (Arabidopsis) (Kao and Nodine, 2019; Zhao et al., 2019). In addition, the vast majority of genes regulating Arabidopsis embryo morphogenesis are zygotically expressed (Nodine and Bartel, 2012; Zhao et al., 2019) and required (Meinke, 2019; Muralla et al., 2011). Therefore, genes are expressed from the zygotic genome during initial stages of embryo development, and the diversification of gene expression programs across plant embryonic cell types contributes to the formation of the basic body plan. Characterizing how gene expression programs are established in individual cell types of early embryos is critical to understand the molecular basis of pattern formation in plant embryos, and more broadly the general and unique principles of embryonic patterning in multicellular organisms.

Forward genetic screens successfully identified many genes required for proper plant embryogenesis (Lukowitz et al., 2004; Mayer et al., 1998; Meinke, 2019), but relatively few mutations in genes encoding cell-specific transcriptional regulators were recovered. This is at least partially due to the high degree of genetic redundancy among plant transcription factors that typically belong to multigene families (Riechmann, 2002). As an alternative approach, RNA populations can be characterized to infer gene-regulatory processes underlying cellular differentiation events. Transcriptomes generated from early embryos at various stages of development have accordingly yielded insights into the biological processes operating during different embryonic phases (Belmonte et al., 2013; Hofmann et al., 2019; Xiang et al., 2011; Zhao et al., 2019). However, these transcriptomes were generated from whole embryos. Additional studies have revealed genes that are preferentially expressed in broad (Belmonte et al., 2013; Casson et al., 2005; Chen et al., 2021; Slane et al., 2014; Zhou et al., 2020) or more specific regions (Palovaara et al., 2017) of plant embryos, but either lack cellular resolution or were contaminated with RNAs derived from the maternal seed coat that encompasses the developing embryo (Schon and Nodine, 2017).

Single-cell mRNA-seq (scRNA-seq) has been instrumental towards understanding developmental events at cellular resolution over the past decade (Chen et al., 2019; Hwang et al., 2018). Several studies have applied these approaches to plant tissues (Brennecke et al., 2013; Efroni and Birnbaum, 2016; Jean-Baptiste et al., 2019; Ryu et al., 2019; Satterlee et al., 2020; Shulse et al., 2019; Song et al., 2020; Xu et al., 2021; Zhang et al., 2019), but scRNA-seq has yet to be reported for individual cell types in plant embryos. This is primarily due to the presence of rigid cell walls that hold plant cells together. Although cell walls can be removed by enzymatic treatment of tissues that are easy to access, such protoplasting techniques remain impractical for early embryos because they are deeply embedded within maternal seed tissues. Single-nucleus mRNA-seq (snRNA-seq) (Habib et al., 2016) offers an alternative method to inspect transcriptomes at single-cell resolution in plants and has been recently applied to endosperm tissues within seeds (Long et al., 2021; Picard et al., 2020). Here, we present a workflow to obtain contamination-free high-quality transcriptomes from individual early embryonic nuclei followed by their assignments to the precursors of the most fundamental plant tissues. Remarkably, these initial embryonic cell types already express characteristic sets of genes, have different evolutionary trajectories and appear to be regulated by distinct epigenetic and transcriptional mechanisms.

## RESULTS

### Acquisition of contamination-free transcriptomes from individual embryonic nuclei

To acquire single-cell transcriptomes of early Arabidopsis embryos, we utilized fluorescence-activated nuclei sorting (FANS) coupled with single-nucleus RNA-seq (snRNA-seq) (Fig. 1A,B). More specifically, we used a transgenic line expressing nuclear-localized green fluorescent protein under the control of the embryo-specific *WUSCHEL-RELATED HOMEOBOX 2* (*WOX2*) promoter (pWOX2::H2B-GFP, pWOX2::tdTomato-LTI6b; hereafter referred to as pWOX2::NLS-GFP) to fluorescently label nuclei in embryos but not the surrounding endosperm or maternal tissues (Fig. 1A) (Gooh et al., 2015). We chose to focus on globular stage embryos because this is when the precursors to the most fundamental plant tissues emerge along apical-basal and radial embryonic axes (Palovaara et al., 2016). Briefly, we fixed siliques or seeds containing globular stage embryos with a low concentration of dithiobis (succinimidyl propionate) (DSP) before nuclei isolation to preserve RNA. Nuclei were also stained with 4′,6-diamidino-2-phenylindole (DAPI), and intact nuclei were selected based on DAPI profiles (Fig. S1A,C). Embryonic nuclei were then isolated based on their strong GFP signal (Fig. S1B,D) and sorted individually into 96-well plates. Fixed nuclei were decrosslinked with dithiothreitol (DTT) to enable the generation of cDNA and the Smart-seq2 protocol (Picelli et al., 2014a; Picelli et al., 2014b) was used to construct next-generation sequencing (NGS) compatible libraries. NGS libraries were then sequenced on an Illumina HiSeq 2500 followed by the alignment of NGS reads to the Araport11 transcriptome (Cheng et al., 2017) and transcript quantification by Kallisto (Bray et al., 2016) (Fig. 1B). After quality controls (see Methods), 534 out of 744 (72%) nuclei were retained for further analyses. A total of 24,591 genes were detected from all nuclei with an average of 440,289 aligned reads and 2,576 detected genes per snRNA-seq library (Fig. S1E,F and Table S1). Therefore, our approach allowed us to acquire high-quality RNA-seq libraries from hundreds of individual embryonic nuclei.

**Figure 1.**
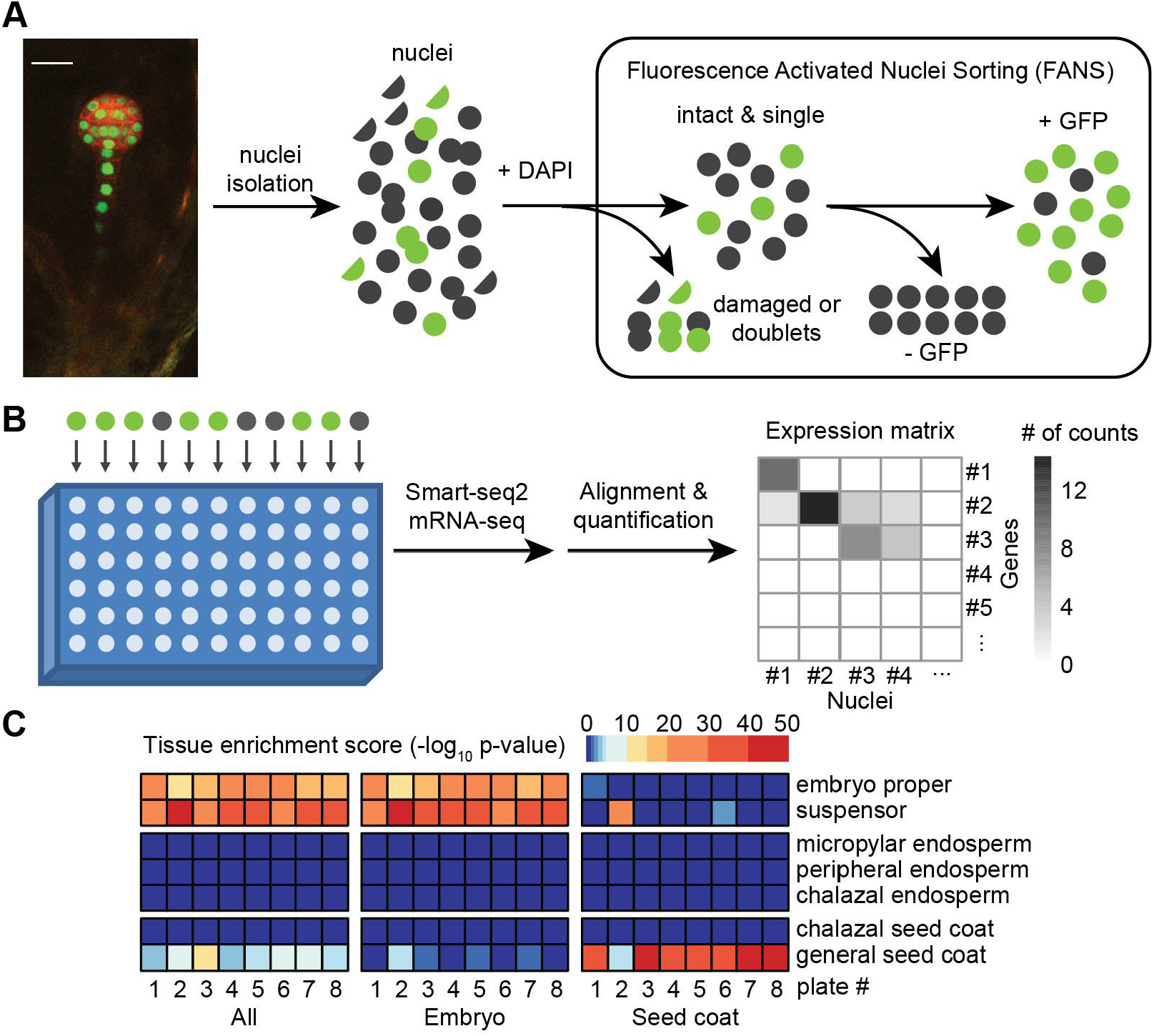
Acquisition of contamination-free transcriptomes from individual embryonic nuclei. (A) Schematic illustration of how single embryonic nuclei were collected. GFP-positive fixed nuclei from pWOX2::NLS-GFP transgenic developing seeds were sorted and collected by FANS (see Methods for details). The scale bar represents 20 μm. (B) Diagram of how single-nucleus libraries were generated with a modified Smart-seq2 protocol (Picelli et al., 2014a; Picelli et al., 2014b), sequenced and individual gene expression quantified (see Methods for details). (C) Maternal contamination assessment and removal. Nuclei from each plate were assigned as embryonic or seed coat-derived according to the unsupervised clustering and tissue enrichment tests (Figs. S2A-C). Tissue enrichment tests based on the mean expression of all nuclei or nuclei categorized as embryo or seed coat are shown.

Contamination of early embryonic mRNA-seq datasets with RNAs from surrounding maternal seed tissues has been a major limitation to embryo transcriptomics (Schon and Nodine, 2017). To evaluate the level of maternal contamination in individual snRNA-seq libraries, we applied the tissue enrichment test (Schon and Nodine, 2017). Although we attempted to achieve 99.9% accuracy with our stringent FANS selection (Fig. S1), embryonic nuclei comprised only 0.1-1% of seed nuclei and thus false positive events were non-negligible and further filtering was required. Accordingly, 20-50% of the snRNA-seq libraries per plate were significantly enriched for either seed coat or endosperm transcripts while remaining snRNA-seq libraries were enriched for embryonic transcripts or had ambiguous identities (Fig. S2A). To systematically identify contaminated snRNA-seq libraries, we conducted unsupervised clustering on all libraries and labeled them according to their tissue enrichment scores (Fig. S2B,C). Because clusters 12 and 13 were enriched for libraries with seed coat contamination, we excluded them from subsequent analyses (Fig. S2B,C). To further evaluate how well we could remove non-embryonic nuclei, we combined the expression levels of snRNA-seq libraries from each plate and performed tissue enrichment tests (Fig. 1C). Retained and excluded nuclei were enriched for embryonic and seed coat cell types, respectively. Moreover, transcriptomes from the retained nuclei were more similar to published embryonic transcriptomes (Hofmann et al., 2019; Nodine and Bartel, 2012) than those from discarded nuclei (Fig. S2C). Altogether, our stringent criteria allowed us to successfully remove non-embryonic snRNA-seq libraries and obtain 486 high-quality snRNA-seq libraries from embryonic cells.

### Identification of embryonic cell types

Because unsupervised uniform manifold approximation and projection (UMAP) clustering of snRNA-seq libraries was able to distinguish embryo proper and suspensor nuclei, but not individual cell types (Fig. S3A), we used enrichments and depletions of known cell-specific transcripts in each nucleus to determine how likely each nucleus was from each cell type. We first identified 174 reference genes expressed in embryos from the literature and recorded their expression patterns as either expressed or not expressed in the cell types present in globular embryos (Table S2, Fig. 2A). More specifically, nine cell types are found in globular embryos (Palovaara et al., 2016) and can be classified based on whether they derive from the larger basal cell or smaller apical cell formed upon zygote division. The corresponding basal cell lineage (BCL) consists of the terminally differentiated suspensor, which connects maternal tissues with the embryo proper, as well as the columella and quiescent center initials that are precursors to distal regions of the root meristem. Unlike the BCL, the apical cell lineage (ACL) divides along the radial embryonic axis to form concentric tissue layers. The outermost protoderm, middle ground tissue initials and innermost vascular initials produce the epidermal, ground and vascular tissues, respectively; whereas the shoot meristem initials will produce aerial tissues after germination. Presence or absence of these reference genes were then used in hypergeometric tests to compute cell-type scores for each nucleus of these 9 cell types in all 486 snRNA-seq libraries (see Methods). We then performed UMAP clustering on the cell-type scores, and identified 12 clusters that were each enriched for a specific cell type (Fig. 2A,B). We also identified one cluster (cluster 4) that was enriched for multiple cell types and had substantially less genes detected per snRNA-seq library compared to the other clusters. We discarded the snRNA-seq libraries belonging to this cluster from subsequent analyses because of their poor quality, which may be due to being generated from aggregated or fragmented nuclei. Cell-specific reference transcripts tended to co-localize to the same cluster (Figs. 2C,S3B) indicating that clustering on cell-type scores recapitulates expression patterns of reference markers. For example, WOX5 and JACKDAW (JKD) transcripts are highly enriched in the quiescent center initials (Haecker et al., 2004; Welch et al., 2007) and co-localize to cluster 12. Therefore, by highlighting the differences among cell types based on a reference gene set we were able to resolve the 486 snRNA-seq libraries into 12 clusters representing distinct cell types.

**Figure 2.**
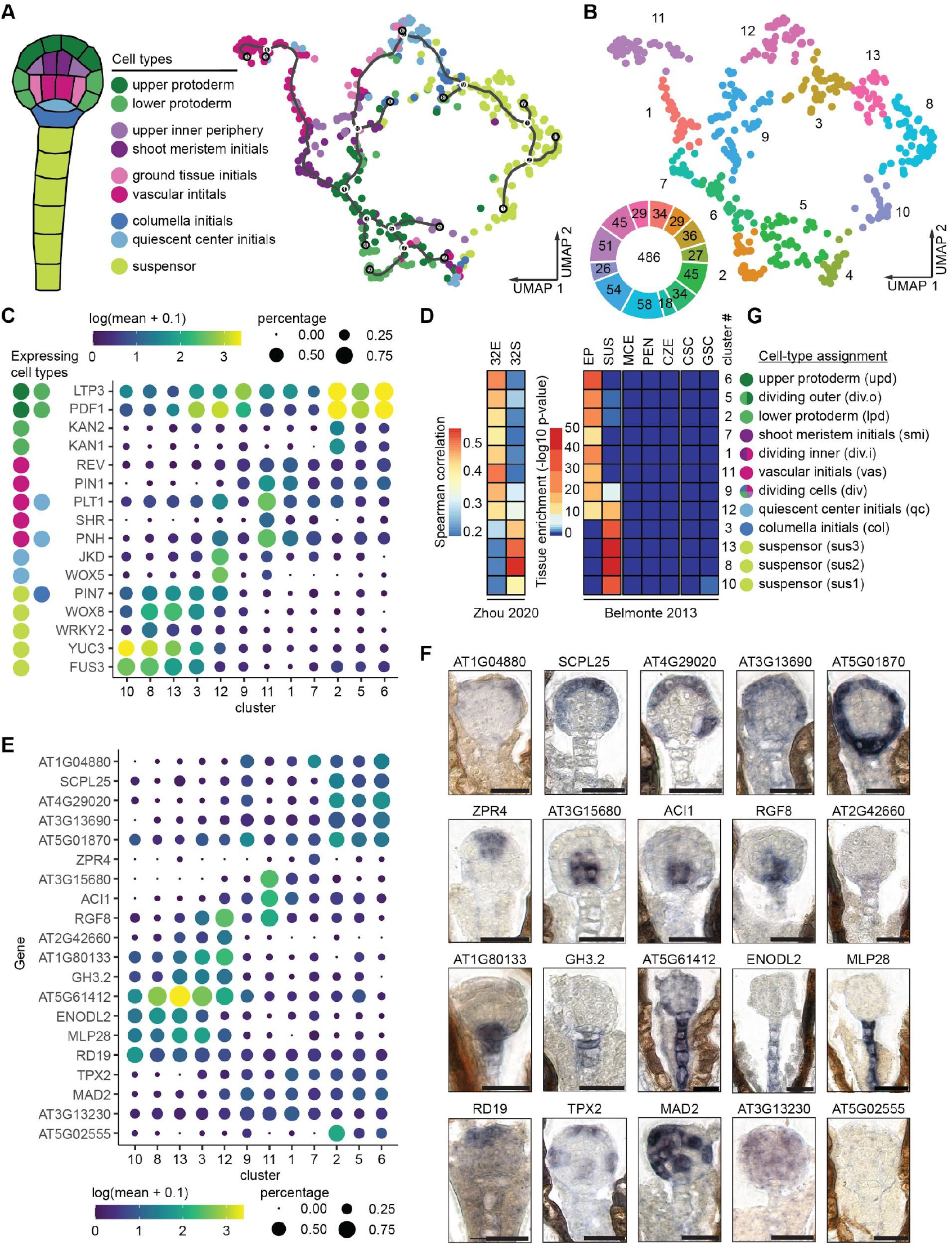
Identification of embryonic cell types. Resolving nine defined cell types by supervised clustering. (A) Marker genes expressed in at least one of the nine cell types were used to calculate cell-type scores with hypergeometric tests. Each dot represents a nucleus and nuclei were labeled according to the cell type with the highest cell-type score. (B) The thirteen clusters corresponding to A. The number of nuclei for each cluster is indicated in the donut plot. (C) Dot plots illustrating expression patterns of known cell type-specific markers. The cell types in which a marker gene is expressed were color-coded according to the diagram in A. The sizes of dots represent the percentage of nuclei in which the transcript was detected for each cluster, and the colors represent the log_10_-transformed mean expression levels of each cluster. (D) Spearman’s correlation coefficients between cluster mean expression and published globular-stage embryo proper (32E) and suspensor (32S) transcriptomes (Zhou et al., 2020) (*left*). Tissue enrichment test results based on cluster mean expression using published transcriptomes from seed tissues as a reference (Belmonte et al., 2013; Schon and Nodine, 2017) (*right*). EP, embryo proper; SUS, suspensor; MCE, micropylar endosperm; PEN, peripheral endosperm; CZE, chalazal endosperm; CSC, chalazal seed coat; GSC, general seed coat. (E) Dot plots of expression patterns of transcripts selected for RNA in situ hybridization (ISH) as in C. Dot plots for the remaining 13 RNA ISH candidates are shown in Fig. S3E. (F) Representative RNA ISH images for 20 selected transcripts. The remaining 13 RNA ISH candidates are presented in Fig. S3F. Scale bars represent 20 μm. Quantification of RNA ISH images is shown in Fig. S4. (G) Assigned cell types and abbreviations for clusters. Asterisks indicated the cluster identities with higher confidence based on in situ and in silico validations.

To independently test these marker-based predictions, we compared the transcriptomes of each cell cluster with published transcriptomes from the embryo proper and suspensor regions of globular embryos (Belmonte et al., 2013). In agreement with the marker-based assignments, clusters 8, 10 and 13 were exclusively enriched for suspensor transcripts based on tissue enrichment tests (Fig. 2D). Also consistent with the cell type assignments, clusters 1, 2, 5, 6, 7 and 11 were enriched for only embryo proper transcripts. Cluster 9 had mixed cell type assignments and accordingly was enriched for both embryo proper and suspensor transcripts. Most of the nuclei in clusters 3 and 12 were respectively labelled as columella initials and quiescent center initials, which are situated between the suspensor and embryo proper. Whereas cluster 3 was only enriched for suspensor transcripts, cluster 12 was enriched for both suspensor and embryo proper transcripts. We also confirmed these results with another published transcriptome dataset generated from embryo propers and suspensors of globular embryos (Zhou et al., 2020) (Fig. 2D). As further support for the cell type assignments of the clusters, three genes not included in our reference list were recently found to be specifically expressed in vascular initials (Smit et al., 2020) and all three were specific to cluster 11 (Fig. S3C) along with other vascular-expressed genes (Fig. 2C).

We then used RNA in situ hybridization (ISH) to further evaluate the marker-based assignments of snRNA-seq clusters to individual cell types. We selected 33 genes without reported expression patterns that represented a specific cell or group of cells based on their expression patterns (Figs. 2E,S3D). We could detect RNA ISH signal in at least 50% of embryos for 26 of the 33 probes tested (78.8%) and compared the RNA ISH and snRNA-seq expression patterns for these in more detail (Figs. 2F,S3E,S3F,S4). *AT3G13690*, *AT4G29020* and *SERINE CARBOXYPEPTIDASE-LIKE 25* (*SCPL25*) were expressed at high levels in clusters 2, 5 and 6, and detected by RNA ISH almost exclusively in the protoderm. *AT5G01870* was also highly expressed in clusters 2, 5 and 6, as well as cluster 12, and was detected in the protoderm and columella initials; whereas *AT1G044880* was expressed in clusters 6 and 9, and detected in the upper protoderm. *AT1G80133*, *AT2G42660*, *AT3G54780* and *GH3.2* were highly expressed in clusters 3 and 12, and RNA ISH signals were detected in the columella and quiescent center initials, as well as throughout the suspensors for GH3.2. *LITTLE ZIPPER 4* (*ZPR4*) was specifically expressed in cluster 7 based on snRNA-seq and detected in the shoot meristem initials by RNA ISH. RESPONSIVE TO DEHYDRATION 19 (RD19) transcripts were also detected by RNA ISH in shoot meristem initials, but were moderately expressed in all clusters. Similarly, *AT4G38370* was expressed throughout the clusters, albeit most strongly in cluster 8, but the RNA ISH signal was stronger in the embryo proper. These two apparent discrepancies between gene expression and RNA localization may be due to differences in post-transcriptional regulation among cell types. *ALCATRAZ-INTERACTING PROTEIN 1* (*ACI1*), *AT3G15680* and *AT3G15720* were expressed most highly in cluster 11 and detected in vascular initials with RNA ISH. *ROOT MERISTEM GROWTH FACTOR 8* (*RGF8*) was highly expressed in clusters 11 and 12, and RGF8 transcripts were detected in vascular and columella initials. *AT5G61412*, *BETA GLUCOSIDASE 17* (*BGLU17*), *COBRA-LIKE PROTEIN 6 PRECURSOR* (*COBL6*), *CYSTEINE ENDOPEPTIDASE 1* (*CEP1*), *EARLY NODULIN-LIKE PROTEIN 2* (*ENODL2*), *MAJOR LATEX PROTEIN 28* (*MLP28*) and *SPERMIDINE DISINAPOYL ACYLTRANSFERASE* (*SDT*) were expressed in clusters 8, 10 or 13, and all were detected in suspensors by RNA ISH. *AT3G13230*, *MITOTIC ARREST-DEFICIENT 2* (*MAD2*) and *TARGETING PROTEIN FOR XKLP2* (*TPX2*) were highly expressed in clusters 1, 5 and/or 9, and corresponding RNA ISH produced ‘salt-and-pepper’ patterns, which are indicative of cell-cycle regulated genes. Accordingly, we observed that clusters 1, 5 and 9 were enriched for mitotic phase regulated transcripts (Menges et al., 2003) (Fig. S3G). Genes preferentially expressed in clusters 1, 5 and 9 also tended to be localized to the subprotoderm, protoderm or both layers, respectively. Therefore, our results suggested that clusters 1, 5 and 9 represent dividing subprotoderm (dividing inner; div.i), protoderm (dividing outer; div.o) and dividing cells in general (div), respectively (Fig. 2G). Altogether, our in silico and in situ validations indicated that we can assign groups of snRNA-seq libraries to the major cell types present in globular embryos: the suspensor (sus1, cluster 10; sus2, cluster 8; sus3, cluster 13); columella initials (col; cluster 3), quiescent center initials (qc; cluster 12); vascular initials (vas; cluster 11); shoot meristem initials (smi; cluster 7); and the lower and upper protoderm (lpd, cluster 2; upd, cluster 6) (Fig. 2G).

### General characteristics of transcriptomes from embryonic cell types

To provide a concise and uniform parameter to examine gene expression patterns across embryonic cell types, we calculated “enrichment scores” in each of the 12 clusters for the 13,893 transcripts detected in ≥10% of nuclei in ≥1 cluster. Enrichment scores are a combination of the deviations of a transcript’s mean levels and the percentage of nuclei it was detected in for each cluster relative to the other 11 clusters (see Methods), and thus concisely summarize the relative abundance of each transcript in each cluster. The 250 genes with the highest enrichment scores (top-ranked 250) from each cluster were considered preferentially expressed genes for that cluster. Enrichment scores of known markers matched their reported expression patterns (Figs. 2C and S3D). For example, 74 of the 118 (62.7%) reference genes were within top-ranked 250 genes of at least one cluster including four that were top-ranked: *PIN-FORMED 1/PIN1* (cluster 1), *KANADI 1/KAN1* (cluster 2), *WOX5* (cluster 12) and *WUSCHEL RELATED HOMEOBOX 8/WOX8* (cluster 13). To gain insights into which biological processes are enriched in each embryonic cell type, we conducted gene ontology (GO) term enrichment analyses on the top-250 ranked genes of each cluster (Fig. 3A and Table S4). Significantly enriched GO terms were identified for the top-ranked 250 genes in the div, vas, div.i, smi, lpd, div.o and upd clusters, but not sus1/2/3, col or qc clusters. The inability to detect enriched terms in these BCL clusters may have been due to the limited annotation of genes specifically expressed in these cell types. Consistent with the div, div.i and div.o clusters representing actively dividing cells, GO terms related to progression through mitotic phases (div and div.o) and microtubules (div.i and div.o) were enriched. GO terms related to body axis specification were also enriched in the top-250 ranked genes of the div.i cluster, as well as the vas cluster. The protoderm clusters (lpd and upd) were both enriched for specification of axis polarity and cutin biosynthesis terms within their top-250 ranked genes. Moreover, the top-250 ranked genes of the lpd and upd clusters could be distinguished from each other by their over-representation of epidermal and cotyledon development GO terms, respectively. The top-250 ranked genes of the smi cluster were enriched for genes involved in DNA replication processes including pre-replicative complex assembly, which is consistent with the smi cluster being depleted for mitosis phase markers (Fig. S3G). Overall, the enriched GO terms were consistent with the assigned cluster identities (Fig. 2G) and indicate that we have classified embryonic cell types with distinct functions.

**Figure 3.**
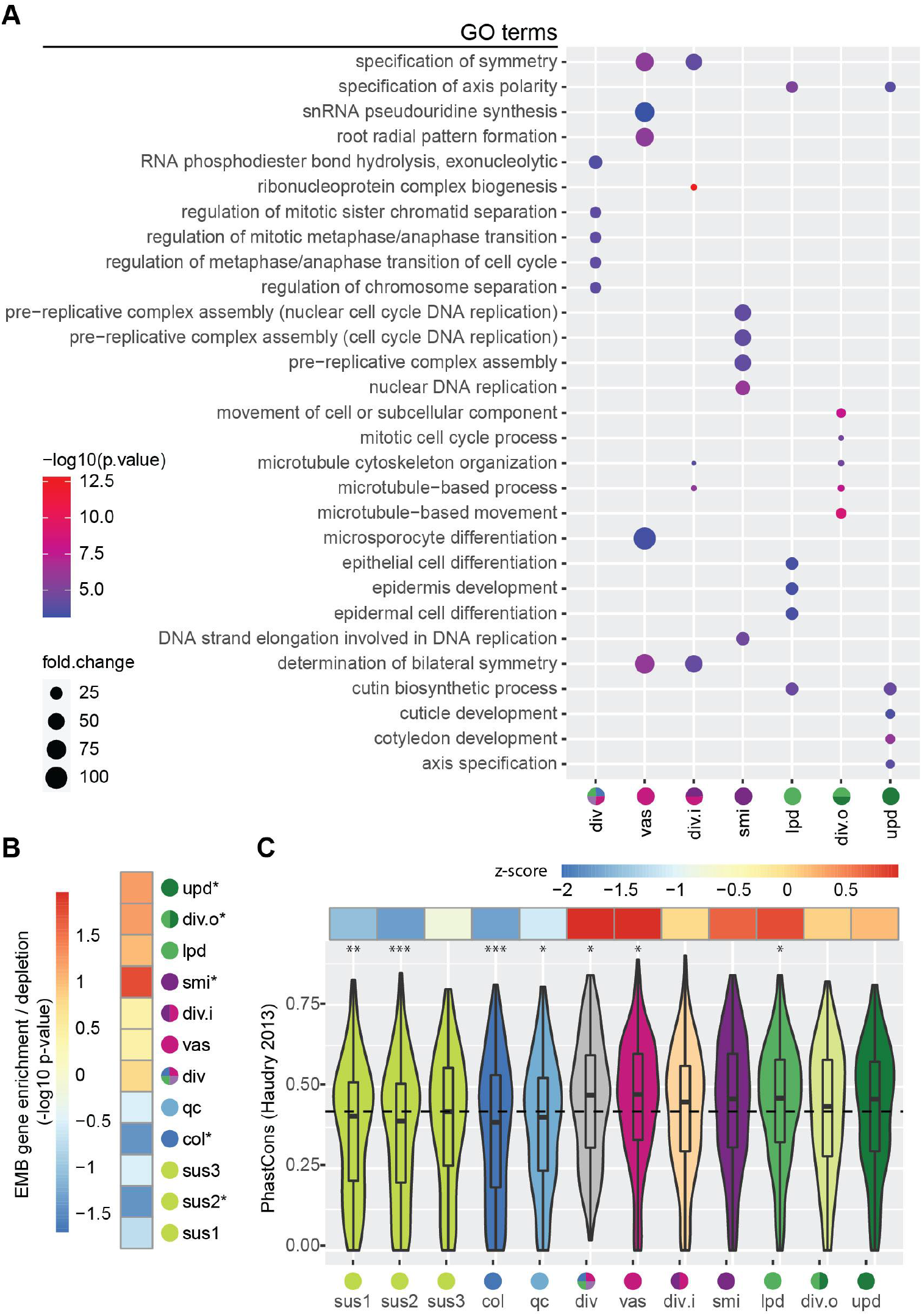
General characteristics of transcriptomes from embryonic cell-types. (A) The top five enriched gene ontology (GO) terms identified by PANTHER for each cluster according to the top-250 ranked genes for each cluster. The suspensor clusters (8,10,13) and hypophysis clusters (3,12) did not have significantly enriched GO terms and thus are not shown. The sizes and colors of the dots represent the fold changes and −log_10_-transformed p-values, respectively. (B) Levels of overrepresentation of embryo-defective (EMB) genes for the top-250 ranked genes for each cluster. Asterisks indicate significant (p ≤ 0.05) enrichment or depletion of EMB genes relative to expectations. (C) PhastCons conservation scores (Haudry et al., 2013) of top-250 ranked genes for each cluster. The mean PhastCons score of all expressed genes is indicated by a dashed line, and deviations from the mean are presented in the upper row as z-scores. The asterisks indicate p-values ≤ 0.05 (*), ≤ 0.01 (**) or ≤ 0.001 (***) based on two-sided Kolmogorov–Smirnov tests with the alternative hypothesis that the cluster conservation score distributions of the top-ranked 250 genes were not equal to that of all expressed genes in embryos. PhastCons and PhyloP scores from another report (Tian et al., 2020) had similar trends (Fig. S5). Cell type abbreviations are as in Fig. 2G.

Next, we tested whether genes essential for embryogenesis are preferentially enriched within the top-250 ranked genes of each cluster. EMBRYO-DEFECTIVE (EMB) genes are a set of genes required for normal embryo development in Arabidopsis (Meinke, 2019). EMB genes were enriched in the top-250 genes of the ACL clusters including significant enrichment in the smi, div.o and upd clusters. By contrast, EMB genes were depleted from top-250 genes of the BCL clusters, including significant depletion in the col and sus2 clusters (Fig. 3B). Further supporting that genes preferentially expressed in the ACL are more likely to be required for proper development than those in the BCL, we found that the top-250 ranked genes within the ACL, and especially the div, vas and lpd clusters, were more highly conserved across Brassicacea species and land plants in general compared to BCL clusters (Haudry et al., 2013; Tian et al., 2020) (Figs. 3C,S5A,S5B). Also consistent with the EMB analyses, the top-250 ranked genes within the BCL clusters were more poorly conserved especially genes enriched in the sus1, sus2, col and qc clusters. Altogether, these results suggested that genes preferentially expressed in ACL clusters, and especially the vas and div clusters are under stronger purifying selection compared to those in BCL clusters, especially the col cluster, which are mutating at a faster rate. This is also consistent with the more variable morphologies of suspensors relative to embryo propers (Chen et al., 2021).

### Transcripts encoding epigenetic regulators vary across embryonic cell types

Soon after fertilization of egg and sperm, epigenetic states are reprogrammed in the new generation (Gehring, 2019). This includes replacement of histones, as well as re-establishment of DNA methylation landscapes genome-wide by small RNA dependent and independent pathways (Bouyer et al., 2017; Ingouff et al., 2010; Jullien et al., 2012; Nagasaki et al., 2007; Papareddy et al., 2020). Because such differential chromatin states can strongly influence gene expression, we examined the transcript levels of genes previously implicated in chromatin regulation. More specifically, we found that 50/191 genes involved in general chromatin features, histone modifications (i.e. acetylation, methylation and ubiquitination), polycomb repressive complexes, DNA methylation or demethylation or small RNA production or activities had enrichment scores ≥2.5 in ≥1 embryonic cell cluster (Erdmann and Picard, 2020; Pikaard and Mittelsten Scheid, 2014) (Fig. 4A). General chromatin factors and components of the polycomb repressive complex tended to vary between the embryo proper and suspensor.

**Figure 4.**
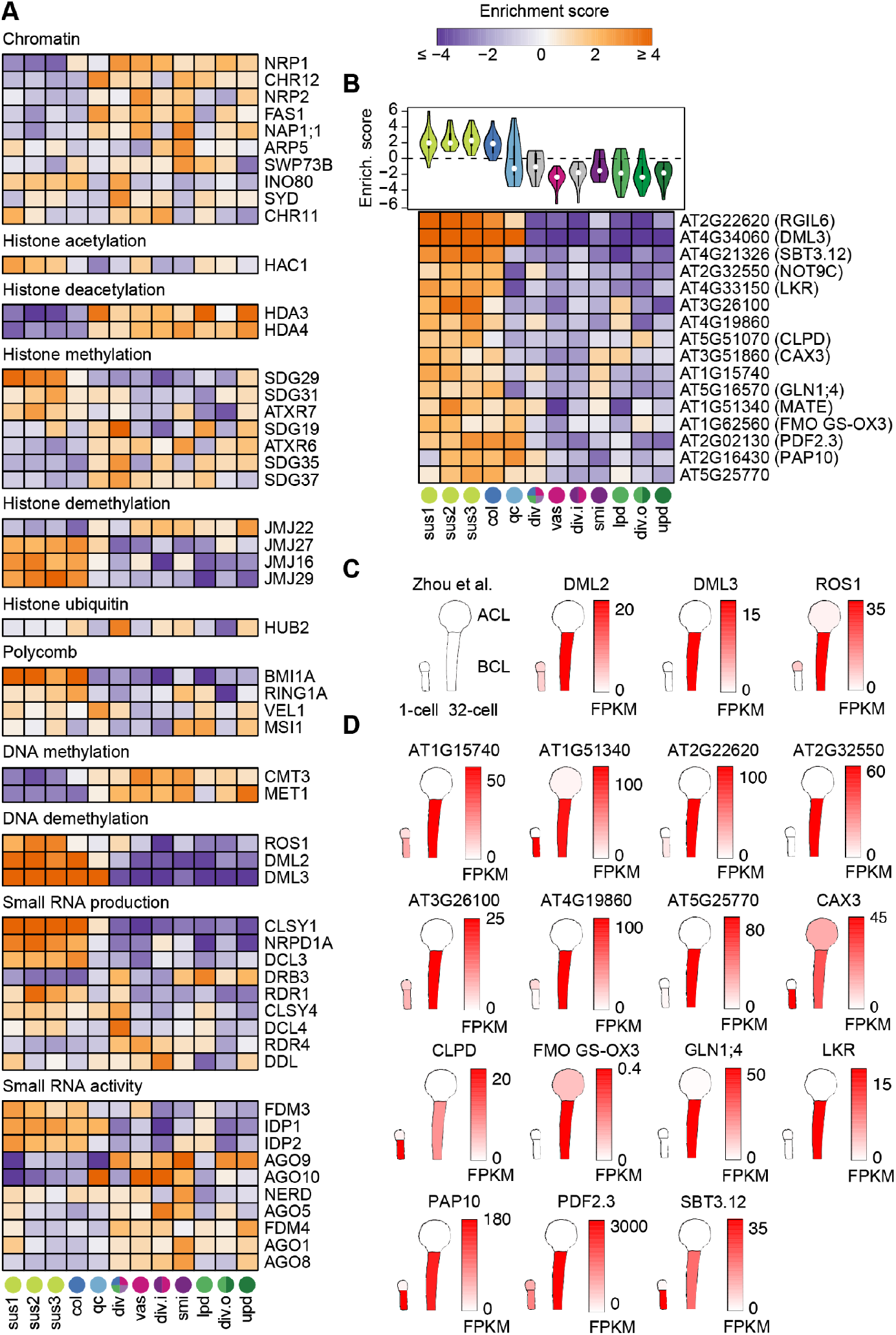
Transcripts encoding epigenetic regulators vary across embryonic cell types. (A) Heatmap illustrating enrichment scores in 12 clusters corresponding to different embryonic cell types. Transcripts with enrichment scores ≥2.5 in ≥1 cell cluster are shown and enrichment scores are colored according to key. Gene names are indicated and cluster identities are marked and color-coded at the bottom according to Fig. 2G. (B) Violin plot (*top*) and heatmap (*bottom*) of enrichment scores for 16/50 ROS1/DML2/DML3 (RDD) targets detected and enriched in the basal cell lineage. (C and D) Schematic representations of RDD transcripts (C) and their putative embryonic targets (D) based on published mRNA-seq from apical and basal cell lineages in 1-cell and 32-cell stage embryos (Zhou et al., 2020). Transcript levels (FPKM; fragments per kilobase of transcript per million mapped reads) are colored according to the keys.

HISTONE ACETYLTRANSFERASE OF THE CBP FAMILY 1 (HAC1) was enriched in the suspensor clusters while HISTONE DEACETYLASE 3/4 (HDA3/4) were enriched in the embryo proper. Moreover, the JUMONJI DOMAIN-CONTAINING16/27/29 (JMJ16/27/29) and JMJ22 histone demethylases were respectively enriched in the suspensor and embryo proper. Interestingly, the terminally differentiated suspensor was enriched for transcripts encoding proteins required for the production of 24-nt small interfering RNAs (siRNAs) such as CLASSY1 (CLSY1), NUCLEAR RNA POLYMERASE D1A (NRPD1A) and DICER-LIKE3 (DCL3). By contrast, genes encoding ARGONAUTE (AGO; AGO1/5/8/9/10) proteins, which bind to small RNAs and mediate gene repression, were enriched in the precursors of the shoot meristem initials. The enrichment of AGOs in shoot meristem initials is supported by previous reports (Gutzat et al., 2020; Jullien et al., 2020; Tucker et al., 2008) and is consistent with small RNA-mediated surveillance pathways that prevent transposon mobilization and other genome de-stabilizing events being enriched in the precursors to all aerial tissues including the gametes. Altogether, these results suggest that small RNA-dependent and -independent pathways establish distinct chromatin environments in individual cell lineage precursors.

The most striking cell-specific enrichments were in pathways affecting cytosine methylation, which is typically associated with transcriptional silencing of transposons and repression of gene promoters (Law and Jacobsen, 2010). CHROMOMETHYLTRANSFERASE3 (CMT3) and METHYLTRANSFERASE1 (MET1) encode DNA methyltransferases that maintain cytosine methylation in the CHG (H ≠ G) and CG contexts, respectively, and both were enriched in the embryo proper. By contrast, transcripts encoding the REPRESSOR OF SILENCING (ROS1), DEMETER-LIKE2 (DML2) and DML3 DNA glycosylases required for the removal of methylated cytosines were highly enriched in the BCL including the suspensor, columella and quiescent center initials. Recently, 275 genes were found to be hypermethylated and downregulated in *ros1 dml2 dml3* triple mutant seedlings undergoing tracheary element differentiation and were considered to be a subset of direct ROS1/DML2/DML3 targets (i.e. RDD targets) (Lin et al., 2020). We detected 50/275 RDD targets in ≥10% of nuclei in ≥1 embryonic cell cluster with enrichment scores ≥2 (Figs. 4B,S6). Sixteen of these RDD targets were highly enriched in the BCL. ROS1, DML2 and DML3 transcripts were increased specifically in the BCL between the 1-cell and 32-cell stages (Zhou et al., 2020) (Fig. 4C). Consistently, most embryonic RDD target candidates were also increased in the BCL during these early embryonic stages (Fig. 4D). Based on these results, we propose that DNA demethylases become activated in the BCL by the globular stage and catalyze the removal of methyl groups from a set of gene promoters to derepress their expression.

### Differential enrichment of transcription factor binding motifs

To gain insights into the transcriptional processes that help define these embryonic cell-specific transcriptomes, we tested whether any consensus DNA motifs from the CIS-BP database of transcription factor (TF) binding experiments (Weirauch et al., 2014) were overrepresentated in the promoters of the top-250 ranked genes of each cluster. A total of 18 TF motif families were overrepresented in at least one cluster (Fig. 5A). WRKY DNA-BINDING PROTEIN 2 (WRKY2) is a transcriptional activator in the BCL and was shown to directly activate WOX8 and WOX9 (Ueda et al., 2011). Consistent with this report, the most overrepresented motif in BCL clusters was the W-box bound by WRKY TFs, and this correlated well with the expression enrichment of WRKY2 (Pearson's r = 0.86; Table S5). In addition to WRKY2, the expression pattern of two other WRKY TFs (WRKY28 and WRKY19) strongly correlated with enrichment of the WRKY motif (Pearson's r = 0.94 and 0.96, respectively). The WOX family binding motif was similarly concentrated in BCL clusters, matching the observed expression pattern of WOX8 and to a lesser extent WOX9. The B3 domain transcription factor *FUSCA3 (FUS3)* is preferentially expressed in the BCL, and the RY motif bound by FUS3 is similarly enriched only in BCL clusters. Maintenance of QC identity in roots requires *JACKDAW* (*JKD*), a member of the INDETERMINATE DOMAIN (IDD) subfamily of C2H2 zinc finger TFs (Welch et al., 2007). The IDD motif is enriched exclusively in the QC initials, where *JKD* is the second highest ranked gene behind *WOX5*. Class IV HD-ZIPs include the L1 layer marker genes *MERISTEM LAYER 1* (*ATML1*) and *PROTODERMAL FACTOR 2* (*PDF2*), and their binding sites are overrepresented in the three protoderm clusters. The binding motifs of R1R2R3 Myb TFs, also known as mitosis-specific activator (MSA) elements, are enriched in the three clusters previously identified as actively dividing tissues (div, div.i, div.o), consistent with the role of R1R2R3 Myb TFs in positively regulating genes required for cytokinesis (Haga et al., 2007).

**Figure 5.**
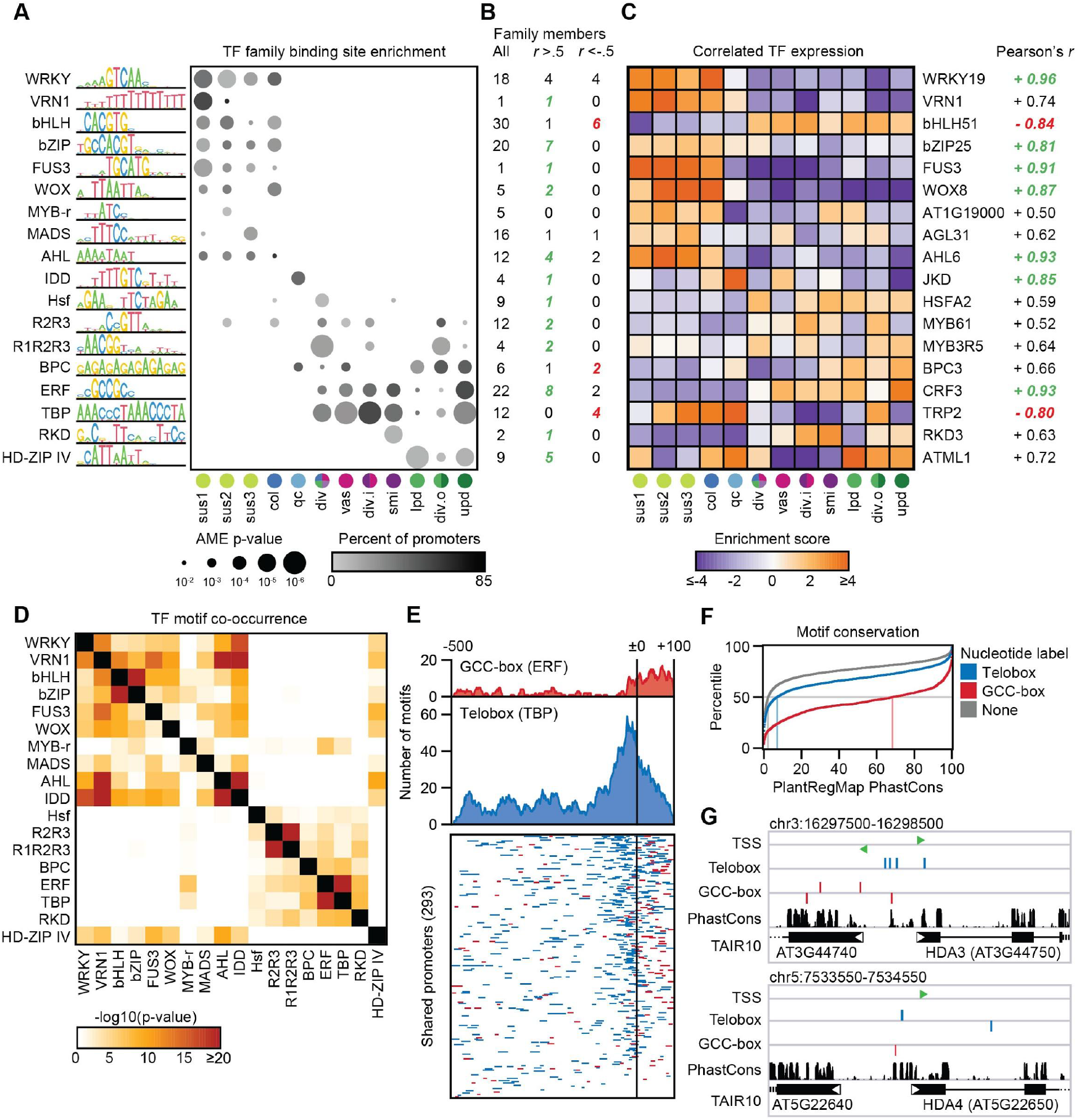
Cluster-enriched transcription factor binding motifs. (A) Dot plot of transcription factor families with DNA binding motifs significantly enriched in at least one cluster. Dot size shows the most significant enrichment (-log_10_ p-value, AME) of a motif in the family; dot color depicts the percentage of the top-250 ranked genes whose promoter contains the specified motif. (B) Number of transcription factors in each family which are detected in the globular atlas (*left*), have a Pearson’s correlation between expression enrichment and motif enrichment across clusters greater than 0.5 (*center*), or less than −0.5 (*right*). (C) Heatmap of expression enrichment scores for the transcription factor within each family whose expression enrichment correlates most strongly to motif enrichment. (D) Pairwise analysis of overrepresentation of a second TF motif co-occurring within the set of top-250 ranked promoters containing a specific motif (p-value, hypergeometric test). (E) Metaplots of the distribution of GCC-box motifs (red) and telobox motifs (blue) relative to the transcription start sites (±0) of 293 top-250 ranked promoters that possess both motifs (*top*). Location of individual motifs in the set of 293 promoters (*bottom*), colored as above. (F) Cumulative frequency of evolutionary conservation score for the promoters in (E), subdivided by whether the nucleotide was contained in a telobox (blue), a GCC-box (red), or neither (gray). Median conservation scores of each group are demarcated by a vertical line. (G) Integrative Genomics Viewer (IGV) browser images of the ±500-bp region around the transcription start sites of HDA3 (*top*) and HDA4 (*bottom*), with putative telobox and GCC-box motifs marked in blue and red, respectively.

To elucidate potential cooperative or competitive relationships between TF binding sites in the globular embryo, the 18 motif families were tested for co-occurrence with each other within the set of promoters from all top-250 ranked genes (Fig. 5D). The major suspensor/embryo proper spatial partition was again observed, with a tendency of BCL and ACL motifs to co-occur in separate sets of promoters. An exception to this pattern was the HD-ZIP IV motif, which was enriched in protoderm clusters, but largely absent from promoters containing other apically-enriched motifs. Instead, HD-ZIP IV motifs tended to occur together in promoters containing motifs characteristic of the BCL, such as FUS3 and WOX binding sites.

One of the strongest pairs of co-occurring motifs was the ETHYLENE RESPONSE FACTOR (ERF) family’s GCC-box and the TELOMERE BINDING PROTEIN (TBP) telobox. In promoters that contain both motifs, the GCC-box tended to occur at or just downstream of the transcription start site, while teloboxes were concentrated immediately upstream (Fig. 5E). For promoters that contained both motifs, nucleotides within a GCC-box or telobox were significantly more conserved in an alignment of 63 land plant species relative to promoter regions that do not include either motif (Haudry et al., 2013; Tian et al., 2020) (Fig. 5F). For example, two histone deacetylases, *HDA3* and *HDA4*, contain one or more conserved copies of the telobox and GCC-box elements in their promoters and are preferentially expressed in all embryo proper tissues relative to suspensors (Figs. 4A,5G). Putative GCC-box binding proteins include the transcriptional activator *DORNRÖSCHEN (DRN)* and four *CYTOKININ RESPONSE FACTORs (CRF1/2/3/10)*, which are expressed in embryo proper tissues enriched for the GCC-box. By contrast, TBP family member transcripts were generally depleted from tissues enriched for TBP binding motifs (Table S5) with the most highly expressed member, *TRP2*, mostly restricted to the BCL. Teloboxes have been reported as functional motifs in Polycomb Responsive Elements (Xiao et al., 2017), and members of the TBP family recruit the Polycomb Repressive Complex PRC2 to teloboxes (Zhao et al., 2018). Therefore, the co-occurrence of teloboxes and GCC-boxes at genes active in the embryo proper is consistent with antagonism between activation by DRN/CRFs and repression by PRC2.

## DISCUSSION

We developed a method to generate high-quality transcriptomes from single embryonic nuclei without detectable contamination from surrounding seed tissues (Fig. 1). Individual nuclear transcriptomes were then grouped according to their cell type, which were validated using published datasets and RNA in situ hybridizations (Fig. 2). This allowed us to construct a gene expression atlas of Arabidopsis embryos at the globular stage when the basic body plan is being established. Our results build upon foundational research examining the divergence of gene expression between the first two sporophytic cell lineages (Belmonte et al., 2013; Chen et al., 2021; Zhou et al., 2020) to help characterize how distinct gene expression programs, and corresponding cell types, are generated during early embryogenesis. Because evolutionary trajectories and transcripts encoding epigenetic factors or transcriptional regulators varied across early embryonic cell types, we surveyed these aspects to gain insights into how distinct gene expression programs are established in early embryos.

Consistent with a recent study, we found that genes preferentially expressed in suspensors tend to diverge more between species compared to those in embryo propers (Geist et al., 2019). Moreover, genes with enriched expression in columella initials were among the most rapidly evolving in the cell types we examined (Figs. 3C,S5). Conflict among siblings for maternal resources is thought to drive adaptive evolution of suspensors (Geist et al., 2019), which support the developing embryo proper and can serve as a conduit for maternally derived molecules (Nagl, 1990; Robert et al., 2018; Shi et al., 2019; Stadler et al., 2005; Yeung, 1980). Because columella initials are situated between the suspensor and embryo proper, they may help regulate communication between mothers and their offspring. Additionally, DNA glycosylases required for demethylation of DNA are up-regulated in the BCL of preglobular embryos, and our results are consistent with them catalyzing the removal of transcriptionally repressive methylation from gene promoters by the globular stage (Fig. 4). Interestingly, genes required for 24-nt siRNA biogenesis (e.g. *CLSY1*, *NRPD1A* and *DCL3*) were preferentially expressed in suspensors whereas transcripts encoding several ARGONAUTE proteins that bind to small RNAs and mediate gene repression were enriched in embryo propers, especially shoot meristem initials from which the gametes are ultimately derived (Fig. 4A). Moreover, small RNAs can move between cells through plasmodesmata (Vatén et al., 2011), which also connect suspensors with embryo propers (Mansfield and Briarty, 1991). Similar to what has been proposed in other terminally differentiated cell-types in reproductive tissues (Calarco et al., 2012; Feng et al., 2013; Hsieh et al., 2009; Ibarra et al., 2012; Mosher and Melnyk, 2010; Slotkin et al., 2009), it is conceivable that suspensors generate large amounts of 24-nt siRNAs that flow into embryo propers and help silence and immobilize transposons to limit their mutagenic potential. Although beyond the scope of the current study, future cell-specific profiling of siRNAs and epigenetic marks in embryos should enable characterization of DNA demethylation and siRNA production in suspensors.

It is well-established that combinations of TFs along body axes drive pattern formation during animal embryogenesis, but relatively little is known regarding combinatorial transcriptional regulation in plant embryos. By examining the relationships between TF expression levels and the enrichments/depletions of their corresponding binding motifs across embryonic cell types, we both verify existing models and provide testable hypotheses for how combinations of TFs integrate positional cues to influence cell-specific gene expression programs in early Arabidopsis embryos (Fig. 6). For instance, we observed characteristic patterns of TF binding motif enrichments and expression patterns in suspensors, quiescent center initials, sub-protoderm, protoderm and shoot meristem initials (Fig. 5). WRKY2 regulates suspensor development by transcriptionally activating WOX8 and WOX9, which in turn are redundantly required for suspensor development (Breuninger et al., 2008; Ueda et al., 2011; Ueda et al., 2017). Accordingly, suspensors were enriched for WRKY and WOX motifs, as well as motifs for FUS3 which has also been implicated in suspensor development (Lotan et al., 1998). Although VRN and AHL TFs do not have reported functions in suspensors, their binding motifs and expression of their corresponding family members (i.e. VRN1, AHL1 and AHL6) were suspensor-enriched, which is consistent with transcriptional regulatory functions. TBP TFs have also not been implicated in suspensor development, and TBP expression and binding motifs were enriched and depleted in suspensor-enriched gene promoters, respectively. TBPs can recruit Polycomb group complexes (PcGs) to target loci and help repress their expression (Zhou et al., 2016; Zhou et al., 2018). Similarly, BASIC PENTACYSTEINE (BPC) TFs can also recruit PcGs to target loci (Hecker et al., 2015; Xiao et al., 2017), and expression of specific BPC family members (e.g. BPC5/7) and BPC binding motifs were enriched and depleted in suspensor-enriched gene promoters, respectively. Because PcGs limit developmental potential of somatic cells (Ikeuchi et al., 2015; Mozgová et al., 2017) and embryo proper gene programs are suppressed in suspensors (Liu et al., 2015; Schwartz et al., 1994), TBPs and BPCs may help silence genes characteristic of the embryo proper in suspensors to preserve their specialized and terminally differentiated cell fates.

**Figure 6.**
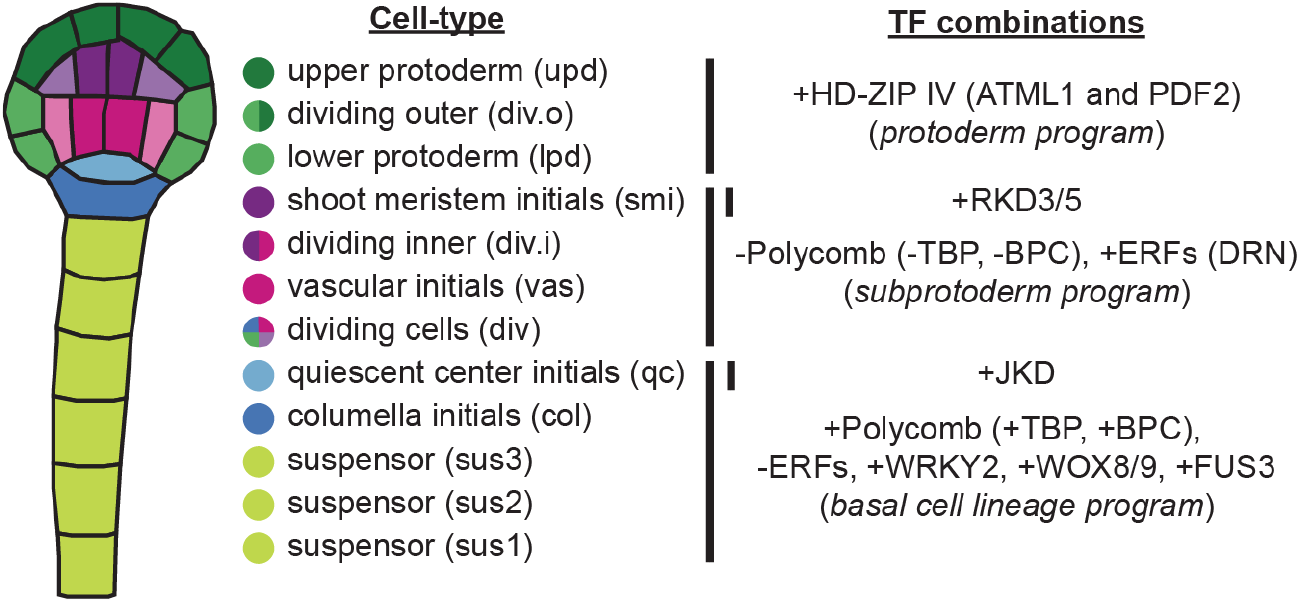
Summary of transcription factor combinations in embryonic cell types. A summary of transcription factor combinations observed or inferred, and which may help establish distinct gene expression programs across cell types (See text for details).

QC initials are derived from the uppermost derivative of the BCL and, unlike suspensors, contribute to post-embryonic tissues. IDD TF binding motifs were specifically enriched in the QC initials and expression of the JKD family member, which is required for QC identity in roots (Welch et al., 2007), was highly enriched in QC initials but not suspensors (Fig. 5). This hints that the superimposition of JKD on the BCL TF combinations could promote QC initial identity in early embryos. Converse to the BCL, the sub-protoderm (i.e. inner cells of the embryo proper) is enriched in co-occurring motifs including the conserved, and often overlapping, TBP and ERF motifs (Fig. 5E,F). Because TBPs are preferentially expressed in the BCL and have been implicated in PcG mediated silencing (Zhou et al., 2016; Zhou et al., 2018), it is possible that in the absence of TBPs, ERFs can bind to these promoters and activate genes that promote developmental potential in the sub-protoderm comprised of stem cell precursors. Among several other ERF family members preferentially expressed in the embryo proper, DRN functions upstream of auxin and binds GCC motifs to promote meristem identity (Chandler et al., 2007; Eklund et al., 2011; Iwase et al., 2017; Kirch et al., 2003). Therefore, we propose that the high degree of developmental potential required for sub-protodermal cells requires a reduction in PcG-mediated silencing and transcriptional activation by ERFs, possibly DRN. The protoderm already expresses genes characteristic of specific processes inherent to the outermost layer during early embryogenesis (Fig. 3A) and is enriched for HD-ZIP IV TF motifs (Fig. 5A). Accordingly, transcripts encoding ATML1 and PDF2 family members were enriched in the protoderm and are required for its specification (Abe et al., 2003; Ogawa et al., 2015). Another cell-specific enrichment of cis-regulatory motifs was observed for RKD TFs in the shoot meristem initials. RKD genes tend to be expressed in reproductive tissues of land plants (Jeong et al., 2011; Koi et al., 2016; Koszegi et al., 2011; Waki et al., 2011) and their over-expression is sufficient to induce expression of undifferentiated cell types (Koszegi et al., 2011; Waki et al., 2011). Therefore, the enrichment of RKD motifs, as well as the preferential expression of RKD3/5 family members, in the shoot meristem initials make RKD3/5 good candidates for future investigation into the establishment of shoot meristem initial gene expression programs.

In addition to providing an early embryonic gene expression atlas, the presented workflow may help guide snRNA-seq experiments on embryos and other plant tissues that are difficult to access. The future application of similar techniques across embryonic stages in Arabidopsis and other species should contribute to a deeper understanding of how gene expression programs are dynamically established during plant embryogenesis. Moreover, integrating snRNA-seq data with other single-cell genomic technologies such as single-cell ATAC-seq (Buenrostro et al., 2015; Cusanovich et al., 2015) may allow further characterization of gene regulatory mechanisms operating in plant embryos. We expect that together with more focussed studies, these genome-wide datasets will accelerate our understanding of the molecular basis of pattern formation in plant embryos.

## MATERIALS AND METHODS

### Plant materials and growth conditions

*Arabidopsis thaliana* accession Columbia (Col-0) plants containing pWOX2::H2B-GFP, pWOX2::tdTomato-RCI2b (pWOX2::NLS-GFP) (Gooh et al., 2015) or no transgenes were grown at 20 to 22°C and 16 h light/8 h dark cycles under incandescent lights (130-150 μmol/m^2^/s) in a climate-controlled growth chamber.

### Nuclei isolation and fluorescence-activated nuclei sorting (FANS)

Developing seeds containing globular embryos from the transgenic pWOX2::H2B-GFP, pWOX2::tdTomato-RCI2b lines and wild type Col-0 were isolated prior to sorting. For each set, developing seeds were isolated with tungsten needles under a stereomicroscope from 20 self-pollinated siliques at stage 17 (Smyth et al., 1990) corresponding to 72 hours after pollination when most embryos are at the early/mid globular stage under the growth conditions used. Developing seeds were isolated at the same time of day to minimize variations caused by circadian rhythms and immediately transferred to 600 μL cooled fixative buffer consisting of 1× Galbraith’s buffer (20 mM MOPS, pH 7.0, 30 mM sodium citrate, 1% Triton X-100, 45 mM MgCl2) and 500 μM dithiobis(succinimidyl propionate) (DSP, ThermoFisher). All buffers used in the nuclei isolation and sorting contained 0.4 U/mL RNAse inhibitor murine (New England Biolabs). Cross-linked samples were incubated with 800 μL quenching buffer (1 M Tris-HCl, pH 7.0, 30 mM sodium citrate, 1% Triton X-100 and 45 mM MgCl_2_) at room temperature for 15 minutes with gentle shaking. The quenched samples were washed twice with 600 μL HG-GB (1× Galbrath’s buffer and 1 M hexylene glycerol, Sigma). The seeds were then gently homogenized with micro-pestles in 1.5 mL microtubes with 200 μL HG-GB. Micro-pestles were rinsed with 400 μL HG-GB, and the homogenized samples were gently pipetted 10 times before incubating at 4℃ for 15 minutes to maximize nuclei release. The partially homogenized samples were then filtered with 30 μm filters and collected in 2 mL microtubes. Another 600 μL HG-GB were added to the 1.5 mL microtube, and filtered and collected through the same 30 μm filter and 2 mL microtube, respectively, to maximize nuclei recovery. The filtered samples were then centrifuged at 1,000 g at 4°C for 10 minutes. The supernatant was carefully removed without disturbing the grayish pellet of nuclei. A fresh aliquot of 1 mL HG-GB and 1 μL of 10 mg/mL DAPI were added into microtubes and the pellet was gently re-suspended. Samples were then washed 5 times, including a 10-minute centrifugation at 1000 g at 4°C and replacement of supernatant with fresh aliquots of 1 mL 1× Galbrath’s buffer. The washed nuclei were then re-suspended in 800 μL 1× Galbraith’s buffer for sorting.

The isolated nuclei were sorted with a BD FACSAria™ III Cell Sorter (BD Biosciences) with a 70 μm nozzle. The scatter gates were adjusted accordingly with Col-0 nuclei. DAPI signals were activated by a 375 nm laser and collected with a 450/40 nm filter. GFP signals were activated by a 488 nm laser and collected with a 530/30 nm filter. To maximize purity, only the droplets containing a DAPI signal within the two peak regions representing 2C and 4C nuclei (Fig. S1A,C) were considered for GFP gating. For GFP gating, a region with low auto-fluorescence and high GFP signal was selected (Fig. S1B,D), which had less than 3 events in Col-0 samples and on average ≥200 events for pWOX2::NLS-GFP samples. Each nucleus passing both DAPI and GFP gating was collected with single-cell settings in 4 μL of cell lysis buffer (Picelli et al., 2014a) supplemented with 25 mM dithiothreitol (DTT) in single wells of 96-well plates.

### Single nucleus RNA-seq (snRNA-seq)

Smart-seq2 libraries were prepared following the published SmartSeq2 single-cell protocol (Picelli et al., 2014a; Picelli et al., 2014b) with an additional 30-minute 37°C incubation before reverse transcription. Libraries were sequenced on an Illumina Hi-Seq 2500 in 50-base single end mode. Sequencing reads from each sample were preprocessed by trimming adapters using cutadapt v2.6 (Martin, 2011) in two steps. First, Nextera adapters (5′-CTGTCTCTTATACACATCTCCGAGCCCACGAGAC-3′) were trimmed from the 3′ end of reads, followed by trimming of template-switching oligos (TSO, 5′-AAGCAGTGGTATCAACGCAGAGTACATGGG-3′) and oligo-dT adapters (5′-AAGCAGTGGTATCAACGCAGAGTACTTTTTTTTTTTTTTTTTTTTTTTTTTTTTT-3′) from the 5′ and 3′ ends of reads, respectively. A Kallisto index was built from a combined FASTA file of all transcript models in EnsemblPlants TAIR10 v40 (https://ftp://ftp.ensemblgenomes.org/pub/plants/release-40/gff3/arabidopsis_thaliana/Arabidopsis_thaliana.TAIR10.40.gff3.gz), 96 ERCC spike-in transcripts and sGFP. Each trimmed sample FASTQ file was pseudoaligned to this index using the command *kallisto quant* with the options *--single--fragment-length 200 --sd 100*.

### Quality control and census count conversion

The TPM table, cell data and gene data were imported into Monocle3 (Cao et al., 2019). Libraries with less than either 100,000 aligned reads or 1,000 detected genes were considered as low quality and excluded from subsequent analyses. The TPM values were then converted to census counts with the census conversion algorithm (Qiu et al., 2017; Trapnell et al., 2014). The census counts were used as gene expression levels in the subsequent analyses.

### Maternal contamination removal and tissue enrichment tests

Gene expression values were used to perform tissue enrichment tests with default settings as described (Schon and Nodine, 2017). The census count expression and metadata of snRNA-seq libraries from 8 plates (Table S1) were constructed as a cell data set (CDS) in Monocle3 with R version 3.6.3. The quality control was done according to Monocle3 guidelines. Genes passing the Monocle3 function detect_genes(CDS, min_expr = 0.1) and expressed in at least 3 nuclei were considered in subsequent analyses. The above quality control steps resulted in a CDS with 534 nuclei and 24,591 genes. An unsupervised Uniform Manifold Approximation and Projection (UMAP) (McInnes et al., 2018) dimension reduction and clustering performed on this CDS resulted in twenty clusters. Two of the clusters (Clusters 12 and 13 in Fig. S2) were dominated by nuclei that resembled the seed coat reference according to tissue enrichment tests, and therefore corresponding nuclei were excluded from subsequent analyses. After contamination removal, a CDS containing 486 globular embryonic nuclei and 23,959 detectable genes was then used for subsequent cell type score calculation and clustering.

### Calculation of cell type scores for globular nuclei and clustering

A set of 174 embryonic marker genes based on either RNA in situ hybridization or transcriptional/translational fusions to fluorescent or beta-glucuronidase (GUS) reporters were collected from the literature (Table S2). Expression levels were recorded as strongly expressed (s), weakly expressed (w), not expressed (n) or non-informative (NA) for each of nine cell types: upper protoderm (upd), lower protoderm (lpd), shoot meristem initials (smi), upper inner periphery (uip), vascular initials (vas), ground tissue initials (grd), quiescent center initials (QC), columella initials (col) and suspensors (sus). The corresponding 174×9 matrix was intersected with expressed genes in our globular snRNA-seq libraries, which had at least one census count in at least 7 nuclei. The resulting 135 expressed marker genes served as the reference for cell type-score calculations, with 56, 52, 38, 43, 62, 43, 56, 51 and 29 positive markers (i.e. strongly or weakly expressed) and 79, 83, 97, 92, 73, 92, 79, 84 and 98 negative markers (i.e. not expressed) for upd, lpd, smi, uip, vas, grd, QC, col and sus, respectively. We utilized two-tailed hypergeometric tests assuming that a nucleus expressing positive and negative markers of a cell type was more or less likely from that cell type, respectively. The resulting p-values were −log_10_-transformed to compute cell type scores. The 486×9 matrix of cell type scores was then used for dimension reduction and clustering. Cluster identities were predicted based on the cell type labels within each cluster.

### Validation of cluster identities

The mean expression values of all nuclei within each cluster were used to perform tissue enrichment tests as previously described (Schon and Nodine, 2017) and to calculate Spearman’s correlation coefficients with published globular stage embryo proper (32E) and suspensor (32S) samples (Zhou et al., 2020). The expression levels of selected markers and three recently reported genes (*PEAR1/AT2G37590*, *DOF6/AT3G45610* and *GATA20/AT2G18380*) (Smit et al., 2020) not included in our reference marker table for tissue score calculation were plotted with Monocle3’s *plot_genes_by_group()* function.

Thirty three RNA ISH candidates without previously reported embryonic expression patterns were selected according to their expression patterns and probe specificity (Table S6). RNA in situ probes were generated from synthesized double-stranded DNA (gBlocks Gene Fragments; Integrative DNA Technologies) and applied as previously described (Nodine et al., 2007). For each probe, 21-122 globular stage embryos (i.e. biological replicates) were examined from 2-8 microscope slides (i.e. technical replicates) for a total of 1,420 embryos from 112 slides (Fig. S4). To minimize potential bias, all in situ images were examined and classified by someone that did not perform the experiments and did not know the identities of the samples.

### Ranked Gene Enrichment

For each cluster of nuclei, a ranked gene enrichment strategy was defined as follows: let *G* be the set of “expressed” genes, defined as all nuclear-encoded and RNA Polymerase II transcribed genes with ≥1 RNA-seq read count in ≥10% of nuclei in ≥1 cluster. For each gene *i* in each nucleus *j*, *CPM_ij_* = 10^6^ × *counts_ij_*/Σ_*g∈G*_*counts_gj_*. Let *C* be a set of nuclei in a cluster and |*C*| the number of nuclei in cluster *C*. Mean CPM of gene *i* in cluster *C* is defined as *μ_iC_* = Σ_*j∈C*_*CPM_ij_*/|*C*|. Proportion detected *p* is defined for each gene *i* in each cluster *C* as the number of nuclei in which gene *i* was detected: 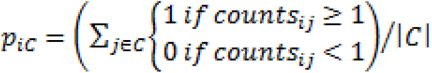. Using one cluster *C* as an ingroup and all other clusters as outgroup *O*, a mean CPM log_2_ fold change of each gene *i* is calculated as 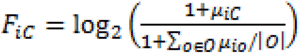, and a mean proportion difference *D_iC_* = *p_iC_* − Σ_*o∈O*_*p_io_*. Both sets *F_C_* and *D_C_* were centered and mean-scaled so that 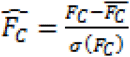, and 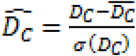, where 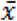 is the mean and the standard deviation. Enrichment magnitude *E_iC_* of gene *i* in cluster *C* is the combined deviation from the mean of *F_C_* and *D_C_*:

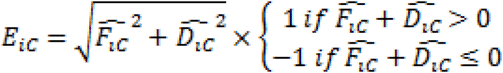

In each cluster, genes were ranked from highest to lowest enrichment magnitude and the first 250 genes and last 250 genes were considered “top-ranked genes” and “bottom-ranked genes”, respectively.

### Gene ontology analyses

The IDs of the top-250 ranked genes for each cluster were submitted to TAIR GO Term enrichment (https://www.arabidopsis.org/tools/go_term_enrichment.jsp) using the PANTHER classification system (Mi et al., 2021) to compute false discovery rates with Fisher’s exact tests. All enriched terms are presented in Table S2. The five most significant GO terms not related to ribosomes are highlighted in Fig. 3.

### Transcription factor binding site analyses

Promoters for all genes were defined as the region 500 bp upstream to 100 bp downstream of the most common 5’ end in nanoPARE datasets of globular stage embryos (Plotnikova et al., 2019). For genes without nanoPARE signal, the most upstream 5’ end annotated in TAIR10 v.46 was used. Transcription factor binding motifs for *Arabidopsis thaliana* were downloaded from CIS-BP (http://cisbp.ccbr.utoronto.ca) (Weirauch et al., 2014). All directly determined motifs were tested for statistical overrepresentation using Analysis of Motif Enrichment (AME, http://meme-suite.org/doc/ame.html) (McLeay and Bailey, 2010) in each cluster by comparing the top-250 ranked gene promoters against a background set of the bottom-250 ranked gene promoters with default parameters. Motifs that were significantly enriched in at least one cluster were collapsed into motif families.

## Contributions

P.K., M.A.S. and M.D.N. conceived the project; P.K. and M.A.S. developed the methodology; P.K. and M.M. performed the experiments; P.K., M.A.S. and M.D.N. prepared figures, wrote and edited the article; M.D.N. acquired funding and supervised the project.

## Acknowledgements

We are grateful to Daisuke Kurihara and Tetsuya Higashiyama for generously sharing pWOX2::H2B-GFP, pWOX2::tdTomato-RCI2b lines. Special thanks to the IMP-IMBA-GMI BioOptics Core Facility, the Plant Sciences Facility and the Next Generation Sequencing Facility at Vienna BioCenter Core Facilities GmbH (VBCF) for their technical support. We also thank Ortrun Mittelsten Scheid, Dolf Weijers and Balaji Enugutti for valuable comments on the manuscript. Ping Kao also personally thanks Kotoha Tanaka and Kaori Sakuramori for their support.

## Competing interests

The authors declare that they have no conflicts of interests.

## Funding

This work was funded by Österreichische Akademie der Wissenschaften (Austrian Academy of Sciences). Research infrastructures in Austria and Czech Republic (RIAT-CZ) project (ATCZ40) funded via Interreg V-A Austria – Czech Republic is gratefully acknowledged for the financial support of the measurements for the preliminary snRNA-seq experiments at the Vienna Biocenter Core Facilities GmbH (VBCF) Next Generation Sequencing Facility.

## Data availability

All sequencing data generated in this study have been submitted to the National Center for Biotechnology Information Gene Expression Omnibus (NCBI GEO, https://www.ncbi.nlm.nih.gov/geo/) under accession number GSE169284.

## Notes

### Competing Interest Statement

The authors have declared no competing interest.

