## Supplementary Information for "Gene Expression Variation in Arabidopsis Embryos at Single-Nucleus Resolution"

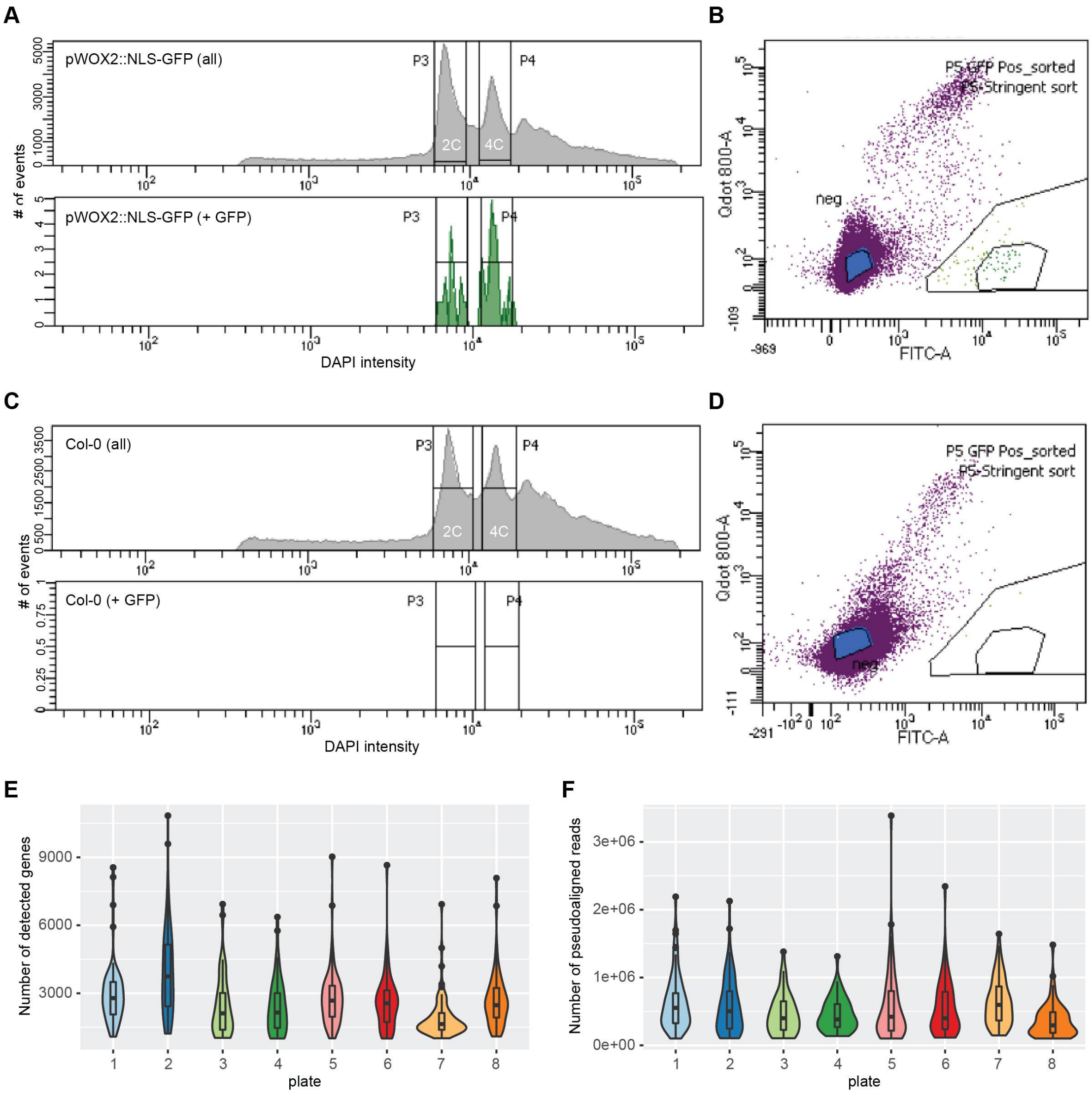


**Figure S1. Embryonic nuclei enrichment with FACS and qualities of snRNA-seq libraries.**

(A and C) Representative DAPI (*top*) and GFP (*bottom*) profiles for the pWOX2::NLS-GFP transgenic line (A) and Col-0 (C). DAPI intensities and the number of events, which include nuclei, are shown on the x and y axes, respectively. Selection gates centered on the 2C and 4C peaks were applied to reduce debris or aggregates.

(B and D) Representative fluorescence scatter plots for the pWOX2::NLS-GFP transgenic line (B) and Col-0 (D), where the x and y axes represent GFP emission and 800-nm auto-fluorescence, respectively. Each dot represents an event (e.g. a nucleus) that passed the DAPI 2C/4C gate. Two regions representing high GFP and low infrared fluorescence were drawn, and the more stringent inner circle was used to enrich GFP-positive nuclei.

(E and F) The number of genes detected (E) and reads aligned to the reference transcriptome (F) in single nuclear libraries from each 96-well plate.

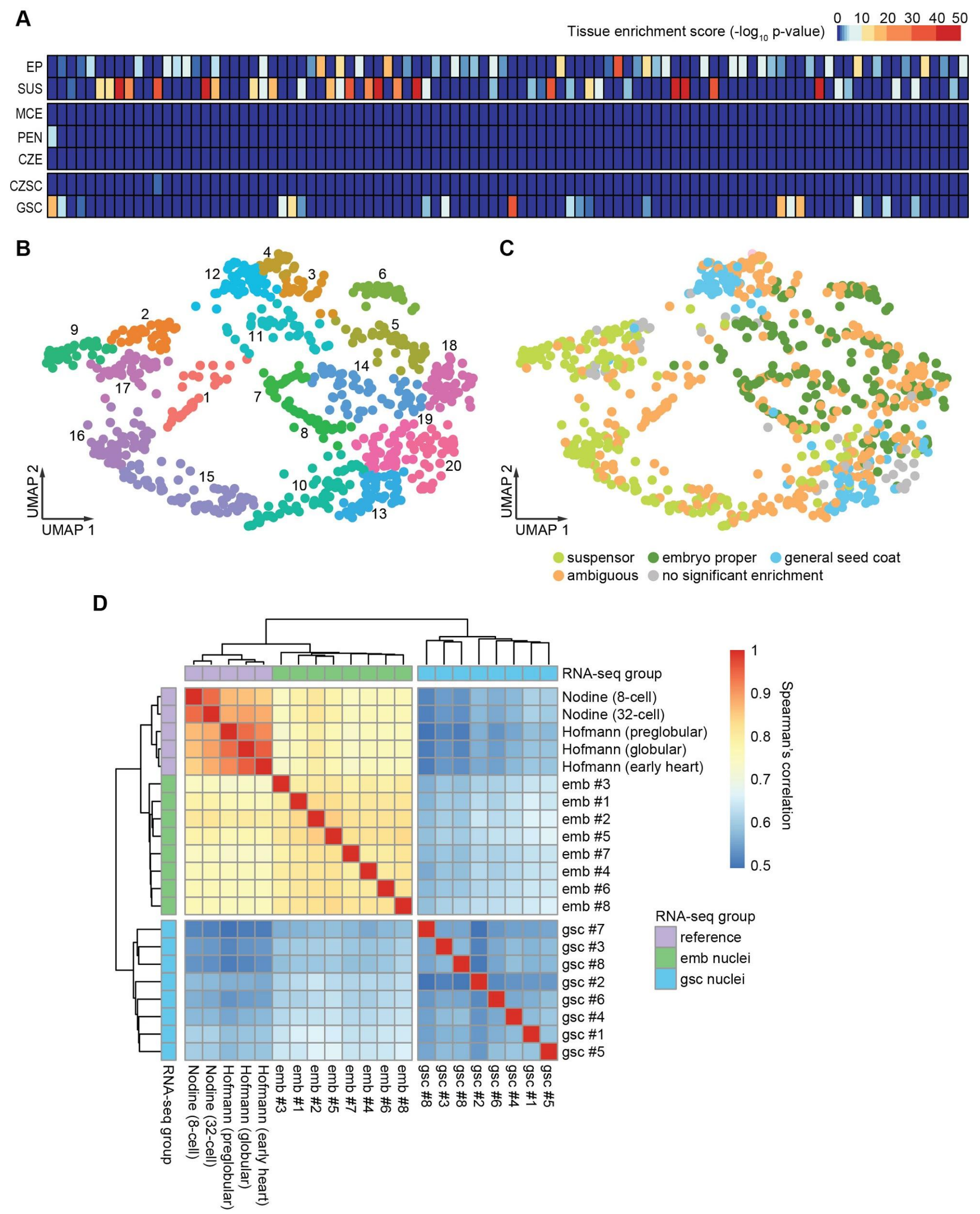


**Figure S2. Assessing and mitigating maternal RNA contamination.**

(A) A representative result of tissue enrichment tests on snRNA-seq libraries from a 96-well plate. Each row represents one of the seed tissue types, and each column represents a snRNA-seq transcriptome. Most nuclei were enriched for one tissue type, indicating that the major source of maternal contamination were false-positive sorted maternal nuclei instead of the ambient RNA. EP, embryo proper; SUS, suspensor; MCE, micropylar endosperm; PEN, peripheral endosperm; CZE, chalazal endosperm; CZSC, chalazal seed coat; GSC, general seed coat.

(**Figure S2**, continued)

(B) Unsupervised clustering and identification of contaminated nuclei. All snRNA-seq libraries with ≥100,000 aligned reads and ≥1,000 expressed genes were clustered and the resulting UMAP plots were color-coded by clusters (*left*) or the significantly enriched tissue type according to tissue enrichment test (*right*). If a snRNA-seq library had no significantly enriched tissue type or was significantly enriched for more than one tissue type, it was labeled as no significant enrichment or ambiguous, respectively.

(C) Heatmap illustrating Spearman's correlation coefficients among the snRNA-seq libraries classified as embryonic (emb) or general seed coat (gsc) and grouped by plate as shown Fig. 1C and published embryonic datasets (Hofmann et al., 2019; Nodine and Bartel, 2012).

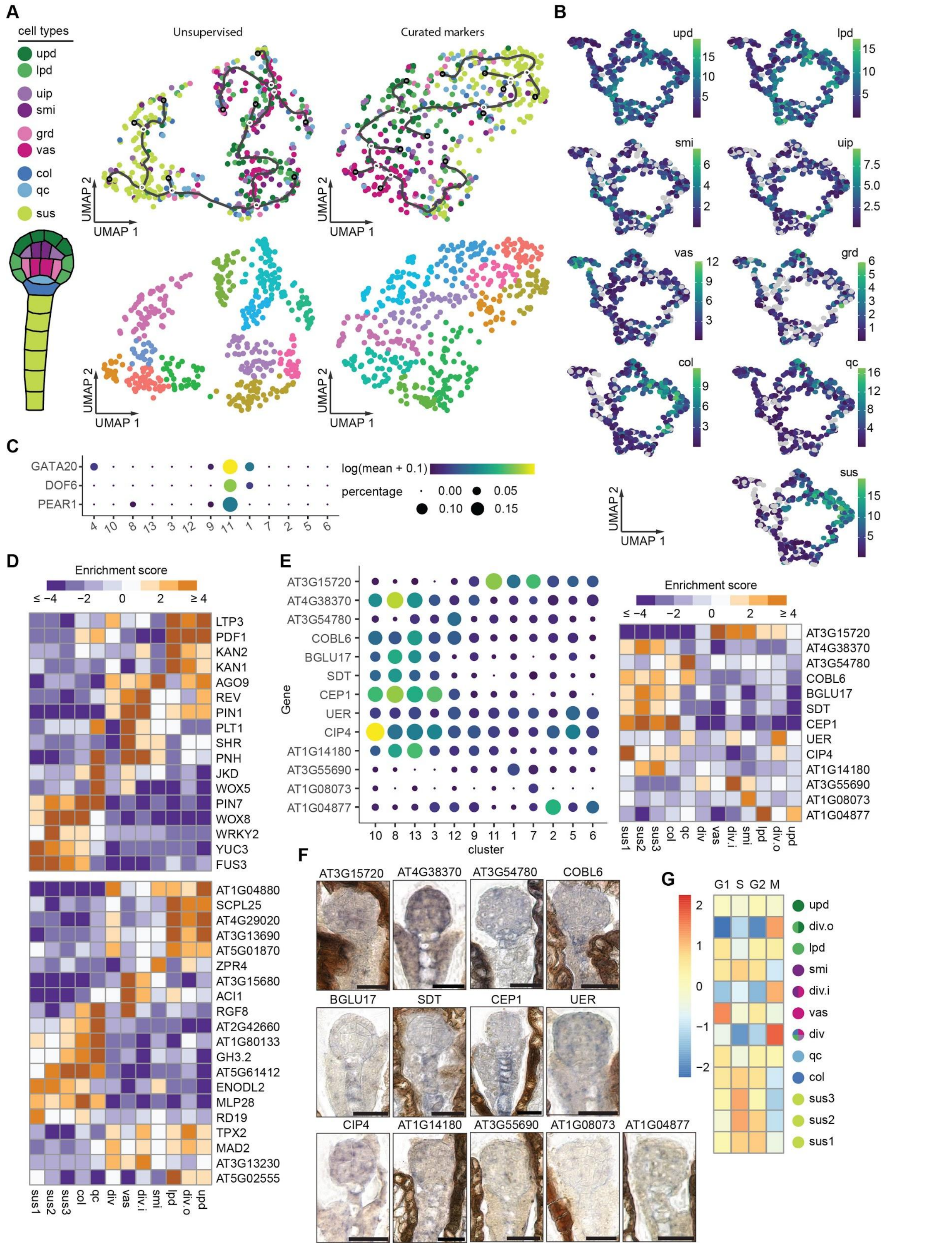

**Figure S3. Resolving cell types by clustering.**

(A) UMAP plots based on unsupervised (*left*) and clustering based on expression of marker genes (*right*). The embryonic cell types are as in Fig. 2A. Each dot represents a nucleus and was colored according to cell type scores (*top*) or cluster (*bottom*).

(**Figure S3**, continued)

(B) Accumulated expression of cell-enriched markers across clusters defined in Fig. 2A and B. Each dot represents a nucleus and was colored according to expression levels (i.e. accumulated census counts) based on the keys. Cell types are indicated in each graph and abbreviations are as in Fig. 2A.

(C) Dot plots illustrating the expression patterns across the clusters defined in Fig. 2A and B for three vascular-specific genes (Smit et al., 2020), which were not included in the marker list used to guide the clustering. The sizes of dots represent the percentage of nuclei the transcript was detected in for each cluster, and the colors represent the $\log_{10}$-transformed mean expression levels of each cluster.

(D) Enrichment scores of the transcripts presented in Fig. 2C (*top*) and Fig. 2E (*bottom*). Cluster identities are indicated at the bottom and are as in Fig. 2G.

(E) Dot plots of expression patterns (*left*) and heatmaps of enrichment scores (*right*) corresponding to the remaining 13 RNA ISH candidates not shown in Fig. 2.

(F) Representative RNA ISH images of the remaining 13 candidates not shown in Fig. 2. Scale bars represent 20 μm.

(G) Enrichments and depletions of cell-cycle related genes among the top-250 ranked genes for each cluster. Cluster identities are as in Fig. 1G.

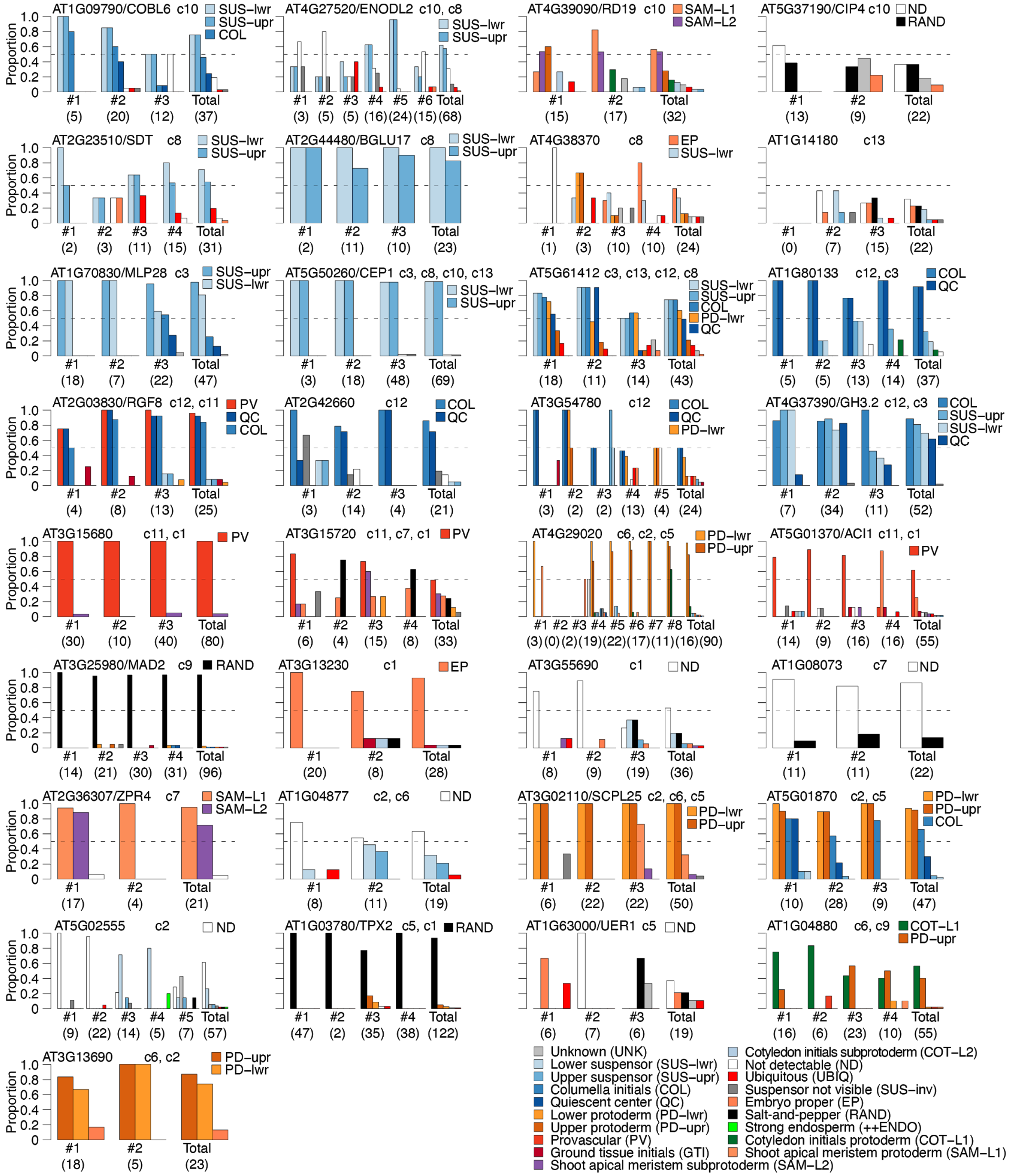


**Figure S4. Quantification of RNA in situ patterns.**

Bar plots illustrating the proportion of globular-stage embryos with RNA in situ signals in various cell types. Each panel summarizes patterns observed for individual transcripts. Corresponding unique gene identifiers (i.e. Arabidopsis Genome Initiative identifiers; AGIs) and, if annotated, common names are shown in the upper-left corner of each panel. Clusters for which transcripts

(**Figure S4**, continued)

are within the top-250 ranked genes are indicated at the top of each graph (e.g. c1, c2, etc.) in ascending order based on their rankings. Proportions of signals are shown for individual slides (i.e. technical replicates), as well as totals for each transcript, and the number of embryos examined (i.e. biological replicates) are noted in parentheses. Legends in each panel are shown for patterns that occurred in >33% of embryos, and the full legend is in the bottom-right corner.

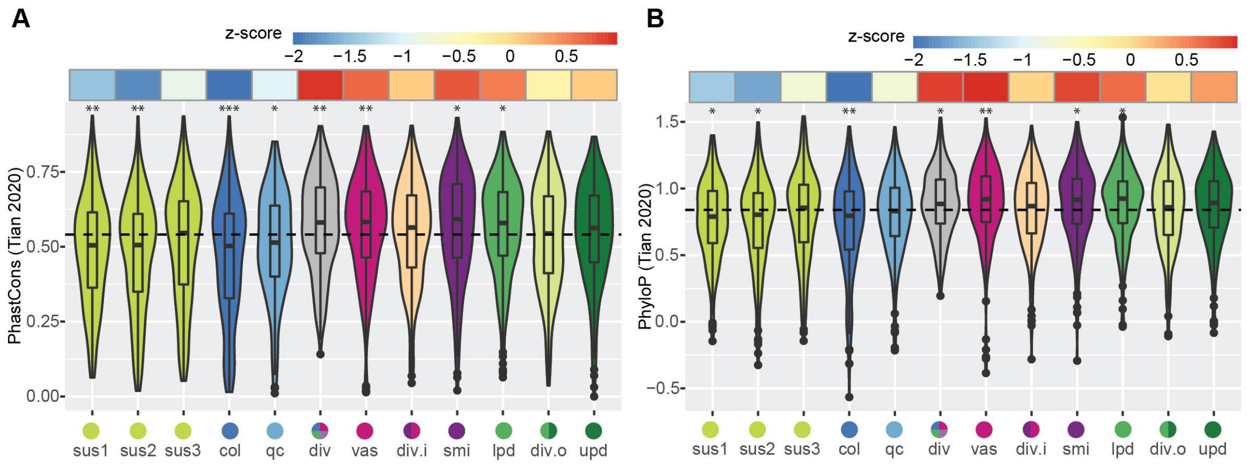


**Figure S5. Conservation score distribution among clusters.**

(A and B) PhastCons (A) and PhyloP (B) conservation scores (Tian et al., 2020) of the top-250 ranked genes for each cluster. The mean scores of all expressed genes are indicated by dashed lines, deviations from the means are presented in the upper row as z-scores. The asterisks indicate p-values ≤ 0.05 (*), ≤ 0.01 (**) or ≤ 0.001 (***) based on two-sided Kolmogorov–Smirnov tests with the alternative hypothesis that the cluster conservation score distributions of the top-ranked 250 genes were not equal to that of all expressed genes in embryos. Cluster identities are as in Fig. 1G.

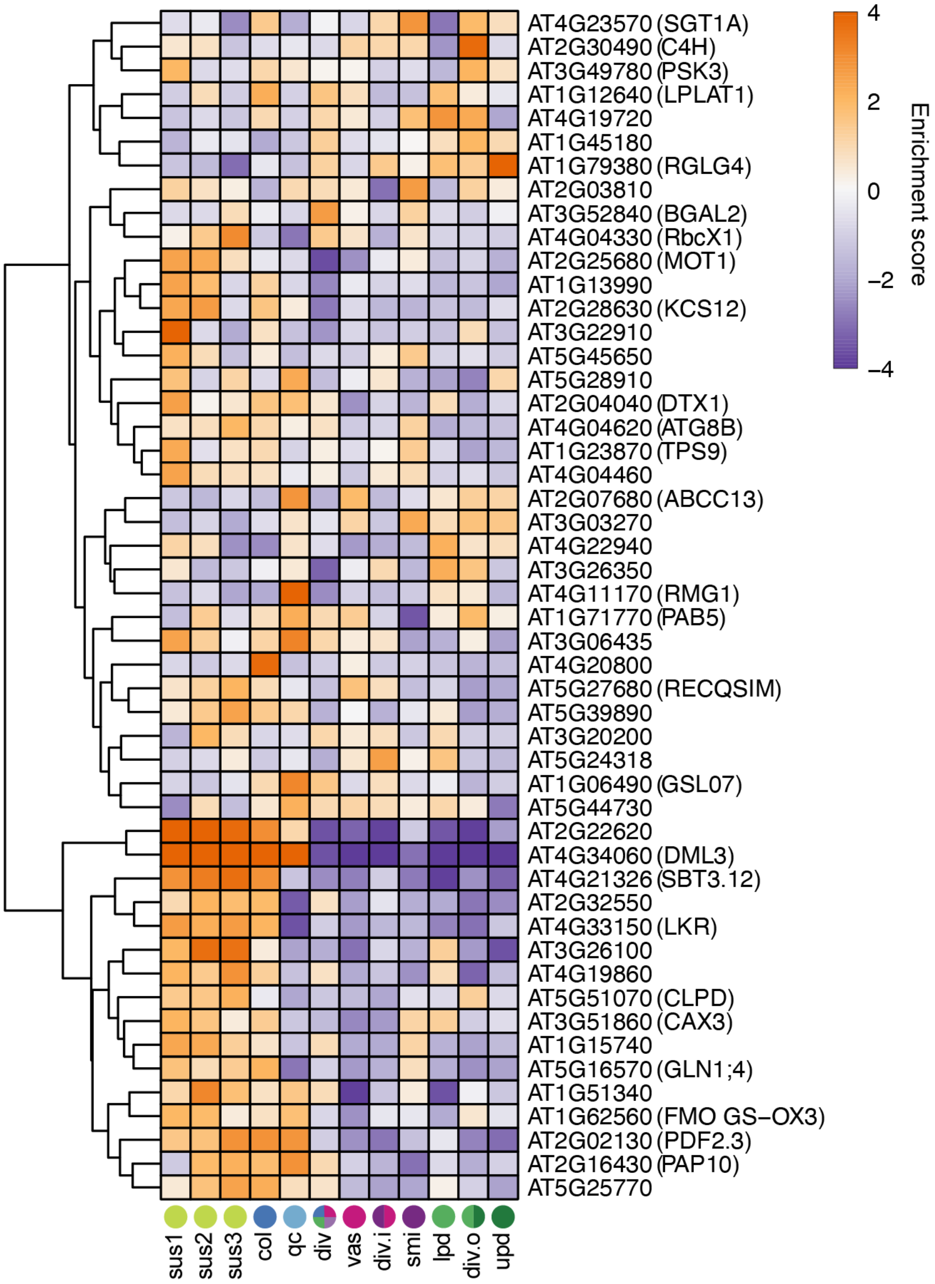


**Figure S6. RDD target candidate expression across embryonic cell types.**

Heatmap of enrichment scores for 50 ROS1/DML2/DML3 (RDD) targets detected in ≥10% nuclei in ≥1 cluster and with enrichment scores ≥2 in ≥1 cluster. Enrichment scores are colored according to the key. Gene names are indicated and cluster identities are marked and color-coded at the bottom. Cluster identities are as in Fig. 1G.

The following supplemental tables are provided separately as Excel spreadsheets:

**Table S1. General information of snRNA-seq libraries and genes detected.**

**Table S2. Curated marker genes used for cell-type score calculation.**

**Table S3. Gene expression data and ranks.**

**Table S4. Gene ontology analyses results.**

**Table S5. Transcription factor motif correlations.**

**Table S6. RNA ISH probe sequences.**
